# Early Natural History of Cardiomyopathy and Cardiac Stress Response in Young Dogs with Golden Retriever Muscular Dystrophy

**DOI:** 10.1101/2024.08.19.608721

**Authors:** Lee-Jae Guo, Peter P. Nghiem, Amanda K. Bettis, Jonathan H. Soslow, Christopher F. Spurney, Joe N. Kornegay

## Abstract

**Background:** Duchenne muscular dystrophy (DMD) and genetically homologous golden retriever muscular dystrophy (GRMD) are X-linked conditions causing progressive muscle wasting and cardiomyopathy. We previously defined a DMD-like dilated cardiomyopathy in adult GRMD dogs. The goal of this study was to extend our work and characterize the early natural history and cardiac stress response in young GRMD dogs.

**Methods:** Age-matched GRMD (N=7), carrier (N=10), and normal (N=8) littermates at 3, 6, and 12 months of age were prospectively enrolled. Electrocardiography (ECG), echocardiography, and cardiac magnetic resonance (CMR) were assessed. To identify early evidence of cardiomyopathy, we conducted a dobutamine stress test in a subset of 17 GRMD and 6 normal dogs. Systolic functions were assessed during dobutamine infusion at 2 months of age, with follow ups (4 GRMD vs. 6 normal) at 4.5 and 6 months.

**Results:** Heart rate and ECG Q/R ratios were greater in GRMD dogs at 12 months, and PR interval was shortened at ≥6 months. There were no differences of echocardiographic systolic function nor circumferential strain in GRMD dogs. CMR left ventricular volumes and myocardial mass were smaller in GRMD dogs ≥6 months, but LGE and T1 mapping did not differ. A diminished inotropic response was seen in GRMD dogs during stress test at 2 months, but not at 4.5 and 6 months.

**Conclusions:** We demonstrated GRMD dogs had a blunted inotropic response at 2 months, and ECG changes and reduced heart sizes ≥6 months. This study substantiated GRMD as a valid animal model for early DMD cardiomyopathy.

**Clinical Perspective:** *What Is New?:*

- This study demonstrates the early natural history of the dystrophin-deficient cardiomyopathy in GRMD model, with early ECG changes and higher heart rate, reduced cardiac chamber sizes, and disproportionally reduced myocardial mass. While with phenotypic variations, imaging markers from ECG, echocardiography, and CMR differentiated the GRMD dogs as early as 6 months of age.
- Systolic function was preserved and no clear evidence of myocardial fibrosis was identified during the subclinical stage of GRMD cardiomyopathy. No cardiac abnormalities were seen in GRMD carriers over the first year of age, aligning with the late-onset cardiomyopathy to carriers.
- A dobutamine stress protocol was established whereby dosages as low as 5 or 10 µg/kg/min intravenous infusion can be used to early differentiate GRMD dogs at 2 months of age. A blunted inotropic response and increased cardiac troponin I to the stress test were found in young GRMD dogs.

*What Are the Clinical Implications?:*

- This study documented the early natural history of GRMD cardiomyopathy, providing a comprehensive dataset and identifying many cardiac imaging markers for preclinical research use. The study further demonstrates the young GRMD dog is a valid animal model for early investigations of DMD cardiac disease.
- The dobutamine stress test and the blunted inotropic response, shown by the changes of fractional shortening (ΔFS) and ejection fraction (ΔEF), provide subclinical markers to detect early cardiomyopathy in very young GRMD dogs.

## Introduction

Duchenne muscular dystrophy (DMD) is an X-linked disease characterized by progressive skeletal muscle wasting and cardiomyopathy that occurs due to out-of-frame mutations in the *DMD* gene with associated loss of the dystrophin protein.^1,2^ Notably, Becker muscular dystrophy patients with in-frame mutations have a mild skeletal muscle phenotype but can develop severe cardiomyopathy later in life.^3^ In this context, improved skeletal muscle function in DMD subsequent to genetic^4^ and cell-based therapies^5^ could potentially increase the cardiac workload.^6^ Indeed, with the advent of more aggressive physical therapy and respiratory care, DMD patients are living longer and more likely to develop cardiomyopathy. The genetically homologous golden retriever muscular dystrophy (GRMD) model has been used extensively to define DMD disease pathogenesis and assess safety and efficacy of proprietary therapies. Most GRMD preclinical studies have been conducted between 3 and 6 months of age, which is analogous to 5-10 years in DMD boys, the time frame of rapid disease progression.^7^ However, while skeletal muscle disease shows dramatic progression in DMD and GRMD during this time, cardiomyopathy has a later onset in both diseases.

Fine et al. published results of electrocardiography (ECG) and echocardiography from a small group of cross-bred dogs with different *DMD* gene mutations over the first year of life.^8^ Previously, our group used advanced imaging modalities including echocardiography and cardiac magnetic resonance (CMR) to show that GRMD dogs typically develop cardiac systolic dysfunction at 30-45 months of age, considerably later than the critical 3–6-month age period.^9^ Additional advanced imaging studies are needed to study cardiac changes in young GRMD dogs. Finding a clinical marker of early cardiac disease would be extremely helpful. While dobutamine stress testing has been used to demonstrate subclinical heart disease in dogs with GRMD and other conditions,^4,5,10–13^ a standard protocol has not been established.

In this study, we comprehensively characterized the early progression of GRMD cardiomyopathy to further facilitate its use in preclinical cardiac trials. First, we compared the natural history of morphologic, hemodynamic, and extracellular myocardial changes in GRMD, carrier, and normal littermates between 3 and 12 months of age. Second, we modified a clinical pharmacologic cardiac stress protocol^14^ for use in young GRMD dogs, with a goal of identifying the ideal dobutamine dosage and the most sensitive outcome parameters.

## Materials and Methods

### Study Animals and Imaging Time Points

All dogs were raised in a colony at Texas A&M University. They were cared for and assessed according to principles outlined in the National Research Council’s Guide for the Care and Use of Laboratory Animals. Studies were approved by the Institutional Animal Care and Use Committee at Texas A&M University. Serum creatine kinase was measured over the first few days of life to identify potential GRMD affected dogs. Genotyping was performed, as described previously,^15,16^ to confirm disease status.

### Age-Matched Natural History Study

This study included 25 dogs from multiple litters (10 GRMD affected, including 5 hemizygous males and 5 homozygous females; 7 GRMD carrier females; and 8 normal males). We evaluated ECG, echocardiography, CMR, and growth rate at 3, 6, and 12 months of age (Figure 1). All imaging data were analyzed using a blinded approach.

**Figure 1.**
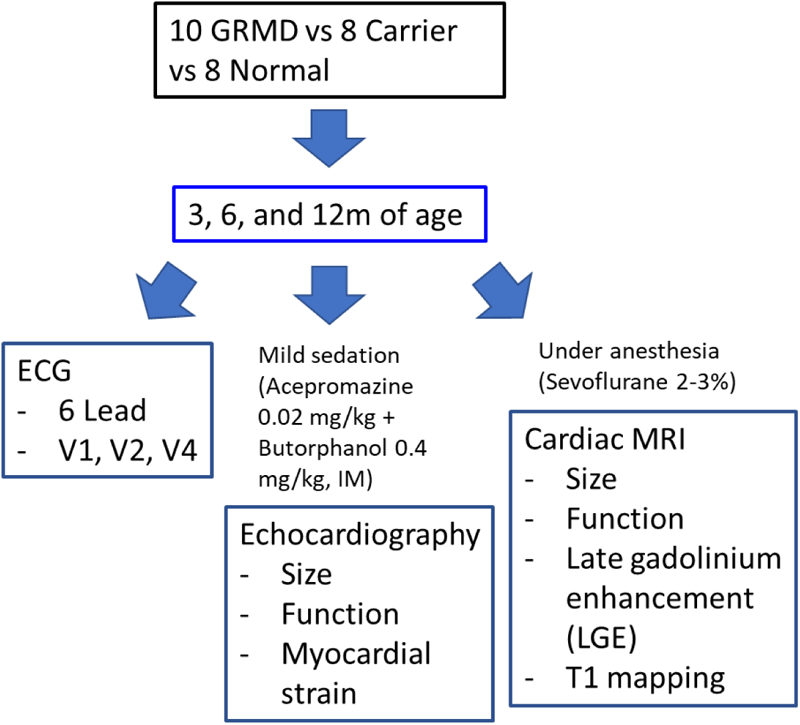
Overview of the time points and imaging modalities in the natural history study

### Electrocardiography (ECG)

Dogs were transported from the kennel and rested in a procedure room at least 10 minutes before the ECG recording. Physical examinations were performed and then the dogs were gently restrained in a right lateral recumbent position. The ECG was recorded using our established protocol without sedation.^17^ The 6-lead, V1, V2, and V4 ECGs were collected on a Philips PageWriter Trim III ECG machine (Philips Medical System, Andover, MA) followed by a 2-minute precordial lead recording using AliveCor Veterinary Heart Monitor (AliveCor, Inc., San Francisco, CA). Heart rate, PR interval, P wave duration and amplitude, QRS duration, and QT interval were measured from lead II. The Q/R ratio was measured on lead II, III, aVF, V2, and V4.

### Echocardiography

Echocardiography was generally performed immediately after the ECG recording. If assessed separately, echocardiography and CMR were collected within 7 days of the ECG. Acepromazine maleate (0.02 mg/kg) and butorphanol tartrate (0.4 mg/kg) were given intramuscularly 10 minutes before the procedure to provide mild sedation and facilitate scanning. The dogs were gently restrained in a lateral recumbent position. A GE Vivid 9 (GE Healthcare, Chicago, IL) ultrasound machine was used as previously described^9^ and images were imported to a GE EchoPAC workstation (GE Healthcare, Chicago, IL) for analysis. Left ventricular (LV) systolic and diastolic function, mitral and aortic flows, average and segmental circumferential strain and strain rate, and myocardial velocity gradient (MVG) were assessed for left heart function. Tricuspid and pulmonary flows were also assessed. The LV end-systolic and end-diastolic volume (LVESV, LVEDV) and ejection fraction (EF) were measured from the right parasternal long-axis view using the modified single-plane Simpson’s rule.^18^ Flows at the mitral and tricuspid valves were obtained from the left apical 4-chamber view. Tissue Doppler imaging (TDI) was used to assess myocardial motion velocities at the mitral annulus from the left apical 4-chamber view. Aortic flow and isovolumic relaxation time (IVRT) were measured from a subxiphoid view. Average and segmental circumferential strain and strain rate^19^ and M-mode analysis^9^ were assessed from a mid-LV short-axis view at the level of the papillary muscles. The MVG was calculated from the differences of endocardial and epicardial velocities using the same short-axis view used for tissue tracking.^20,21^ All echocardiographic parameters were calculated from an average of at least 3 to 5 cardiac cycles.

### Cardiac Magnetic Resonance Imaging (CMR)

The CMR scans were performed immediately after echocardiography. Atropine sulfate (0.04 mg/kg) was given intramuscularly while the dogs were under mild sedation. The dogs were masked with 4-5 % sevoflurane to allow intubation, and then maintained under anesthesia with 2-3 % sevoflurane.

The CMR scans were performed on a 3-Tesla MRI machine (Siemens 3T Magnetom Verio, Siemens Medical Solutions, Erlangen, Germany) with dogs in a dorsal recumbent position. A cardiac dedicated surface coil was used with an ECG or pulse-oximetry gating system. Breath holds or a free breathing sequence were applied, depending on the animal’s breathing pattern and respiration rate. The CMR scan was performed using a previously described protocol,^9^ including assessment of LV function, aortic flow, LGE, and myocardial T1 mapping. Aortic flow was measured using the flow velocity sequence at the aortic valve and then stroke volume (SV) and cardiac output (CO) were measured. LGE imaging was performed 10 minutes after injecting 0.2 mmol/kg gadolinium intravenously. The modified Look-Locker inversion recovery (MOLLI) sequence was used to obtain pre-and post-contrast myocardial T1 maps (Figure 2), followed by measurement of myocardial pre-(native T1) and post-contrast T1 relaxation times for ECV calculation.^22,23^ Blood was collected immediately after the CMR scan to determine the hematocrit value for ECV calculation expressed as a percentage using this formula:

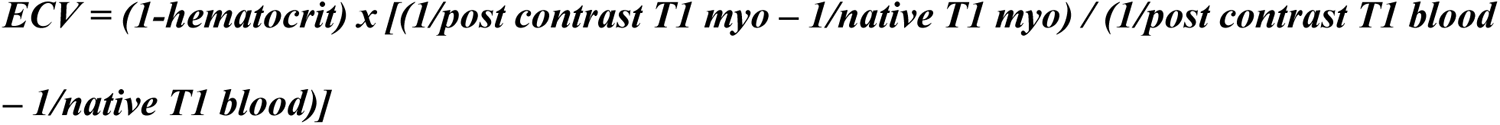

**Figure 2.**
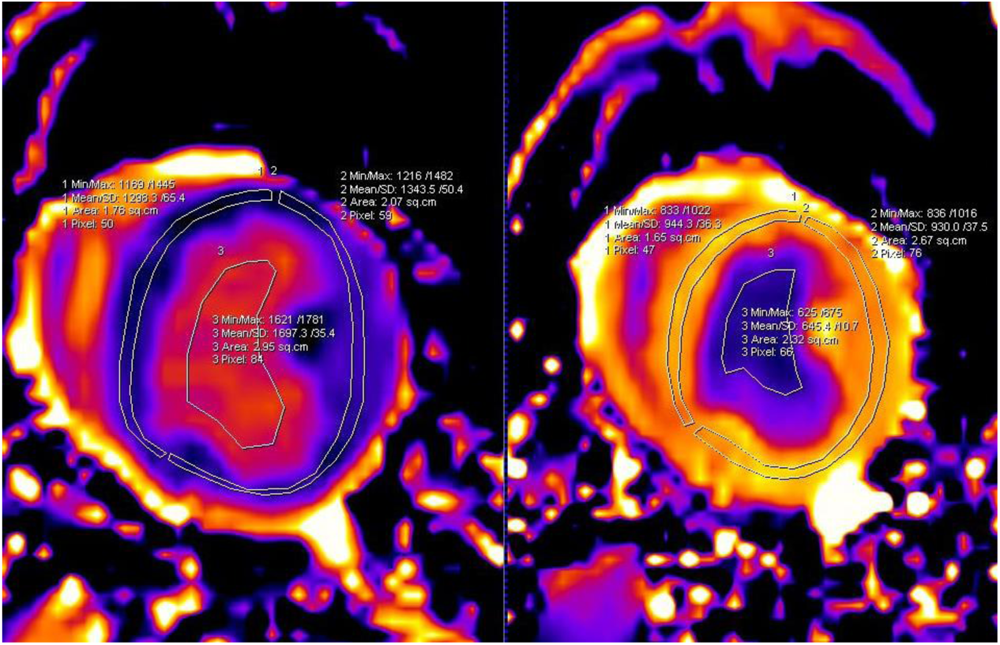
Pre- and post-contrast myocardial T1 relaxation time maps at the mid-LV level. T1 relaxation times were measured from pre- (left) and post-(right) gadolinium myocardial T1 maps at the LV septum, lateral wall, and blood pool. Using hematocrit and the formula of extracellular volume fraction (ECV), ECV values of LV septum and lateral wall were obtained. The pre-contrast myocardial T1 map was used for measurements of native T1 relaxation time. Min/Max, Minimum/Maximum of T1 relaxation time. Mean/SD, Mean/Standard Deviation of T1 relaxation time. Area, area size of the region of interest.

All CMR images were imported into a Siemens Syngo workstation (Siemens Medical Solutions USA, Inc., Malvern, PA) for analysis. Siemens Argus software (Siemens Medical Solutions USA, Inc., Malvern, PA) was used for LV function analysis. Regions of interest were manually drawn by following the endocardial and epicardial edges in all short-axis views in the end-diastolic and end-systolic stages to assess LV function. To obtain more accurate values of the pre- and post-contrast myocardial T1 relaxation time maps, regions of interest were manually drawn by selecting the middle 1/3 of the wall thickness.

### Pharmacological cardiac stress study

An additional subset of 23 dogs from multiple litters (17 GRMD, including 8 hemizygous males and 9 homozygous females; 6 normal males) were assessed using stress echocardiography and stress CMR at 2 months of age. Thirteen of the GRMD dogs were subsequently included in preclinical treatment studies at 3 months so were not subsequently evaluated, leaving 4 GRMD (3 hemizygous males and 1 homozygous females) and 6 normal dogs for evaluation at 4.5 and 6 months of age.

A standardized anesthetic protocol was used for echocardiography, as detailed above.^9^ Dobutamine was administered through a cephalic vein catheter, starting at 5 μg/kg/min for 10 mins. Imaging data were recorded initially at 10 minutes post infusion while the dobutamine was continuously running. Subsequent dobutamine dosages of 10, 15, and 20 μg/kg/min were given, and data were collected after a 5-minute infusion at each dosage (Figure 3). For stress echocardiography, a maximum dobutamine dosage of 25 μg/kg/min was achieved and the dogs were then recovered from anesthesia. The stress echocardiography was performed using an upgraded GE Vivid E9 ultrasound machine (GE Healthcare, Chicago, IL). Right parasternal long-axis 4 chamber and LV short-axis papillary muscle views were recorded at each dobutamine dosage, and the cine images were later analyzed at the workstation for EF and FS measurements. The lead II ECG on the ultrasound machine was used to determine heart rate and monitor cardiac arrhythmia during the stress test.

The stress CMR scans were delayed by at least 24 hours, allowing the dogs to fully recover from stress echocardiography. Dogs were induced and maintained under general anesthesia using the protocol above. The same dobutamine stress protocol was applied during the stress CMR (Figure 3), with a maximum dobutamine dosage of 20 μg/kg/min. The short-axis cine series was serially collected for each dobutamine dosage to obtain LV function data. Heart rate was recorded from ECG gating signal and EF was later analyzed. When all imaging data had been collected, dobutamine infusion was stopped and the dog was recovered from anesthesia.

**Figure 3.**
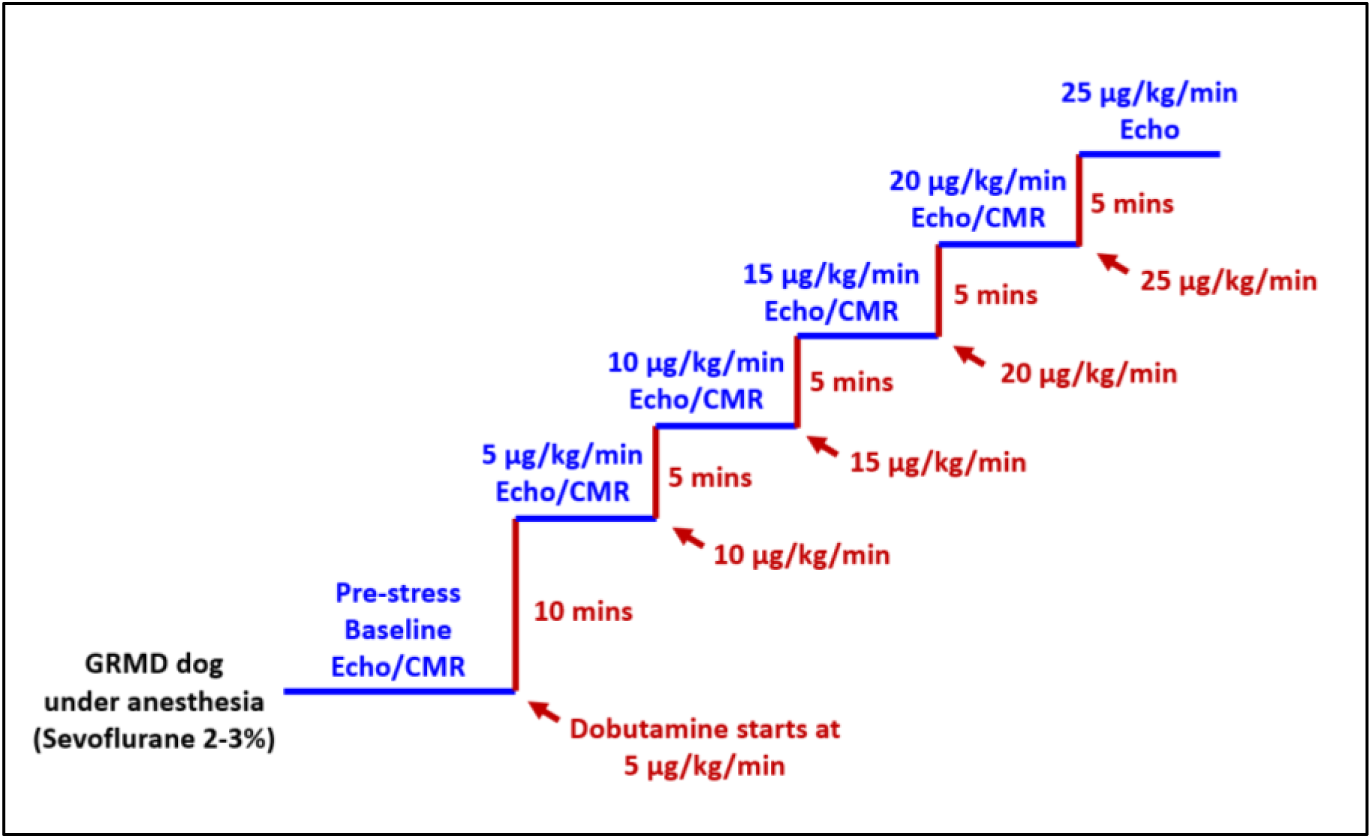
Dobutamine stress protocol used in young GRMD and normal dogs. In our study, the peak dosage was 25 µg/kg/min dobutamine for echocardiography (echo) and 20 µg/kg/min for CMR. The echocardiographic and CMR scans were performed at least 24 hours apart.

To further evaluate the stress effect of dobutamine on the heart, serum cardiac troponin I (cTnI) was measured in 8 affected (1 male and 7 females) and 5 normal male dogs at 2 months of age at the pre-stress echocardiography baseline and post-stress 4 ± 1 hours after the dobutamine peak dosage (25 μg/kg/min). The ultra-sensitive cTnI was measured using the ADVIA Centaur TnI-Ultra assay (Siemens Healthcare Diagnostics Pty Ltd, Bayswater Victoria, Australia), a three-site sandwich immunoassay using direct chemiluminometric technology.

### Statistical Analysis

Statistical analysis was performed using JMP Pro 14-16 software. In the natural history portion of the study, the Wilcoxon rank-sum test was used to compare hemizygous male vs. homozygous female dogs. The Kruskal-Wallis test was used to compare relevant values of affected, carrier, and normal dogs at each time point. If significance was found in the Kruskal-Wallis test, the Steel-Dwass test was used to identify groups that differed significantly. In the cardiac stress study, the Wilcoxon rank-sum test was used to compare the echocardiographic and CMR imaging parameters and cTnI results between affected and normal groups. Wilcoxon signed rank test was used to compare the pre- and post-stress cTnI results for each genotype. *P* value < 0.05 was considered statistically significant in both studies.

## Results

### Animals in Natural History Study

Twenty-five dogs, including 10 affected (5 male and 5 female), 7 carrier (all female), and 8 normal (all male), were assessed for each of the 3-, 6- and 12-month time points.

### Gender Comparison: GRMD Hemizygous Male vs. Homozygous Female

No difference was found between body weights and body surface area (BSA)^24^ of GRMD hemizygous male and homozygous female dogs (n = 5 for each) (Table 1), at 3, 6, and 12 months of age (*P* > 0.05 for all). Only two differences with unclear clinical consequences were found in these same dogs on all ECG, echocardiography, and CMR results. LA/Ao ratios on echocardiography were higher in hemizygous males versus homozygous females at 3 months (*P* < 0.05). The ECG PR interval of hemizygous males was shorter at 12 months (*P* < 0.05) but heart rate did not differ. Therefore, we combined hemizygous male and homozygous female affected dogs as a single group and compared with female carriers and normal male dogs.

**Table 1.** Body weight and body surface area of GRMD affected hemizygous male and homozygous female dogs.

|  | 3 Months |  | 6 Months |  | 12 Months |  |
| --- | --- | --- | --- | --- | --- | --- |
|  | Affected Male | Affected Female | Affected Male | Affected Female | Affected Male | Affected Female |
|  | Median | Median | Median | Median | Median | Median |
|  | (Min–Max) | (Min–Max) | (Min–Max) | (Min–Max) | (Min–Max) | (Min–Max) |
|  | N = 5 | N = 5 | N = 5 | N = 5 | N = 5 | N = 5 |
| Body Weight (kg) | 6.8<br>(4.7–9.2) | 6.0<br>(5.5–8.5) | 11.1<br>(9.2–12.9) | 11.1<br>(9.5–12.9) | 13.3<br>(11.2–16.5) | 14.4<br>(13.3–16.7) |
| Body Surface Area (m<sup>2</sup>) | 0.36<br>(0.28–0.44) | 0.33<br>(0.31–0.42) | 0.50<br>(0.44–0.56) | 0.50<br>(0.45–0.56) | 0.57<br>(0.51–0.65) | 0.60<br>(0.57–0.66) |

### Genotype Comparison of Body Size

Carriers had higher body weight and BSA values than normal (*P* = 0.04) and affected (*P* = 0.02) dogs at 3 months of age (Tables 2 and S1). At 6 and 12 months, body weight and BSA values were lower in affected dogs compared to carriers and normal dogs (P < 0.01 for all). Body weight and BSA values were also lower in carriers versus normal dogs (P < 0.01) at 12 months, likely reflecting a gender effect.

### GRMD vs. Normal Dog Comparison

#### ECG Findings and Heart Rate

Heart rate measured from lead II did not differ at 3 and 6 months of age (Tables 2 and S1). By 12 months, the affected dogs had higher heart rates than normal dogs (*P* < 0.05). The PR interval of the affected dogs was shorter at 6 and 12 months of age, compared to normal dogs (*P* < 0.05 for both). P wave duration and amplitude, QRS duration and QT interval did not differ at any of the three ages. We further compared the Q/R ratio in leads II, III, aVF, V2, and V4 at each of the three time points. The Q/R ratio was higher in affected vs. normal dogs at 12 months of age in leads II, III, and aVF (*P* < 0.05 for all).

While the dogs were under sedation during echocardiography (Table 3), the heart rate measured from M-mode was higher in affected versus normal dogs at 12 months of age (*P* = 0.004). Anesthetized affected dogs also had higher heart rates during CMR (Table 4), at both 6 (*P =* 0.014) and 12 months (*P =* 0.002) of age.

**Table 2.**
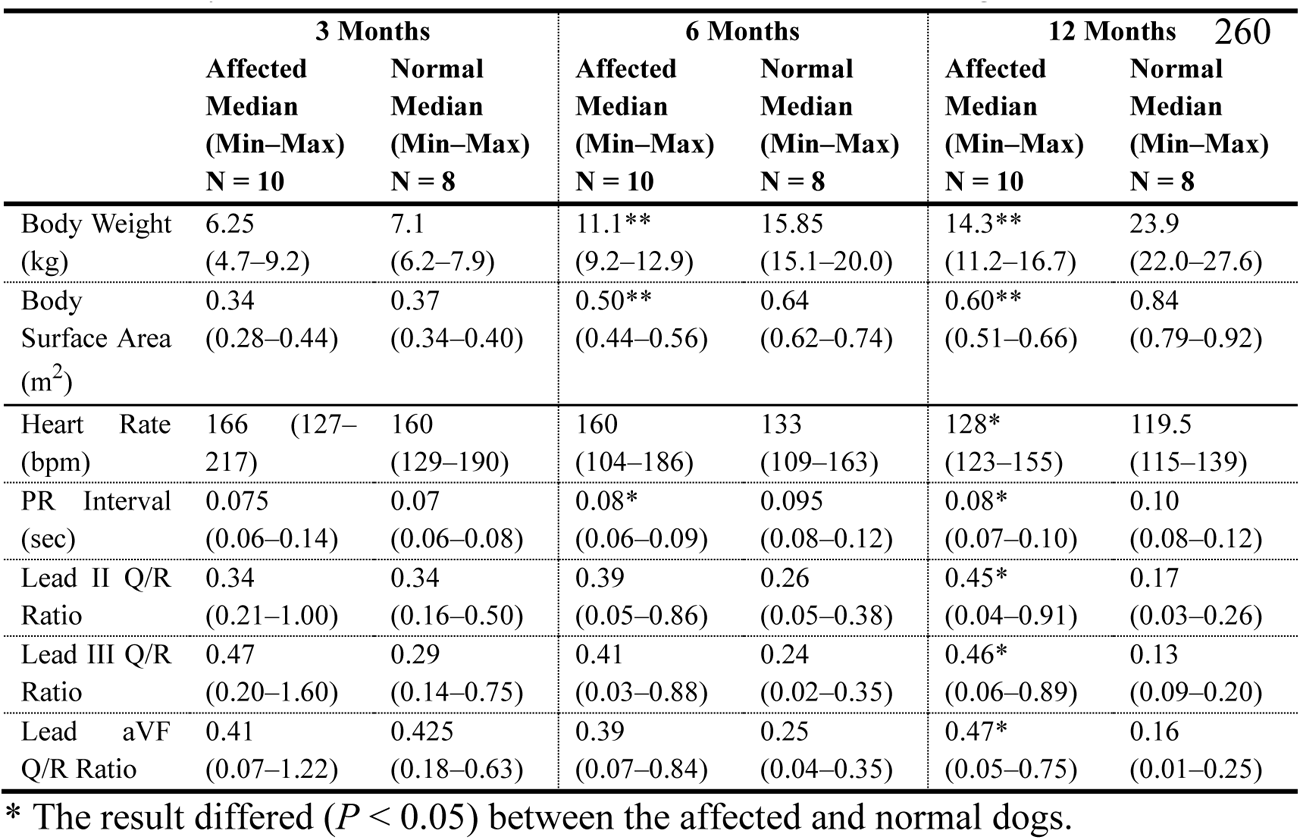
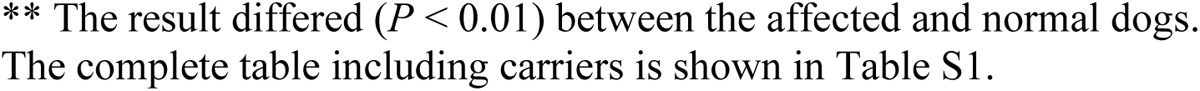
Body size and ECG results from GRMD vs. normal dogs.

|  | 3 Months |  | 6 Months |  | 12 Months |  |
| --- | --- | --- | --- | --- | --- | --- |
|  | Affected<br>Median<br>(Min–Max)<br>N = 10 | Normal<br>Median<br>(Min–Max)<br>N = 8 | Affected<br>Median<br>(Min–Max)<br>N = 10 | Normal<br>Median<br>(Min–Max)<br>N = 8 | Affected<br>Median<br>(Min–Max)<br>N = 10 | Normal<br>Median<br>(Min–Max)<br>N = 8 |
| Body Weight (kg) | 6.25<br>(4.7–9.2) | 7.1<br>(6.2–7.9) | 11.1**<br>(9.2–12.9) | 15.85<br>(15.1–20.0) | 14.3**<br>(11.2–16.7) | 23.9<br>(22.0–27.6) |
| Body Surface Area (m<sup>2</sup>) | 0.34<br>(0.28–0.44) | 0.37<br>(0.34–0.40) | 0.50**<br>(0.44–0.56) | 0.64<br>(0.62–0.74) | 0.60**<br>(0.51–0.66) | 0.84<br>(0.79–0.92) |
| Heart Rate (bpm) | 166 (127–217) | 160 (129–190) | 160 (104–186) | 133 (109–163) | 128* (123–155) | 119.5 (115–139) |
| PR Interval (sec) | 0.075 (0.06–0.14) | 0.07 (0.06–0.08) | 0.08* (0.06–0.09) | 0.095 (0.08–0.12) | 0.08* (0.07–0.10) | 0.10 (0.08–0.12) |
| Lead II Q/R Ratio | 0.34 (0.21–1.00) | 0.34 (0.16–0.50) | 0.39 (0.05–0.86) | 0.26 (0.05–0.38) | 0.45* (0.04–0.91) | 0.17 (0.03–0.26) |
| Lead III Q/R Ratio | 0.47 (0.20–1.60) | 0.29 (0.14–0.75) | 0.41 (0.03–0.88) | 0.24 (0.02–0.35) | 0.46* (0.06–0.89) | 0.13 (0.09–0.20) |
| Lead aVF Q/R Ratio | 0.41 (0.07–1.22) | 0.425 (0.18–0.63) | 0.39 (0.07–0.84) | 0.25 (0.04–0.35) | 0.47* (0.05–0.75) | 0.16 (0.01–0.25) |

\* The result differed ( $P < 0.05$ ) between the affected and normal dogs.
\*\* The result differed ( $P < 0.01$ ) between the affected and normal dogs. The complete table including carriers is shown in Table S1.

#### Heart Size Comparison

We showed that GRMD hearts were smaller than normal on 2D echocardiography (Tables 3 and S2). On M-mode, their left ventricular internal diameters in diastole (LVIDd) and systole (LVIDs) were smaller at 6 and 12 months of age (*P* < 0.01 for all). Affected dogs also had lower interventricular septum thickness in diastole (IVSd) and systole (IVSs) (*P* < 0.01 for both) and left ventricular posterior wall thickness in diastole (LVPWd) and systole (LVPWs) at 12 months (*P* < 0.05 for both). The LV end-diastolic volume (LVEDV) and end-systolic volume (LVESV) measured from the modified Simpson’s rule were smaller in affected dogs than normal dogs at 6 and 12 months (*P* < 0.01 for most). The diameter of left atrium (LA) was smaller in affected dogs than normal dogs at 6 (*P* = 0.02) and 12 months (*P* = 0.001). Additionally, affected dogs had smaller ascending aorta (Ao) diameters at 6 months and lower LA/Ao ratios at 12 months (*P* < 0.05 for both).

Consistent with the 2D echocardiographic findings, affected dogs had smaller hearts compared to normal dogs at 6 and 12 months on CMR imaging (Table 4 and S3), evidenced by lower LVEDV and LVESV and less myocardial mass (Myo Mass) (*P* < 0.01 for most). Strikingly, affected dog’s myocardial mass was two times smaller than normal dogs by 12 months of age (Table 4), while the differences of body weight and body surface area were less (Table 2). A similar but less marked pattern was also shown in the affected dogs at 6 months of age. We further compared the ratios of Myo Mass/Body Weight and Myo Mass/BSA between genotypes. The affected dogs showed lower values of Myo Mass/BSA than normal dogs at 6 months (*P* = 0.004) and lower values of Myo Mass/Body Weight and Myo Mass/BSA at 12 months (*P* < 0.003 for both) (Table S3).

**Table 3.** Echocardiographic results from GRMD and normal dogs.

|  | 3 Months |  | 6 Months |  | 12 Months |  |
| --- | --- | --- | --- | --- | --- | --- |
|  | Affected<br>Median<br>(Min–Max)<br>N = 10 | Normal<br>Median<br>(Min–Max)<br>N = 8 | Affected<br>Median<br>(Min–Max)<br>N = 10 | Normal<br>Median<br>(Min–Max)<br>N = 8 | Affected<br>Median<br>(Min–Max)<br>N = 10 | Normal<br>Median<br>(Min–Max)<br>N = 8 |
| **2D** |  |  |  |  |  |  |
| LVEDV (mL) | 12<br>(7–20) | 12.5<br>(9–18) | 16**<br>(13–22) | 31<br>(26–46) | 20**<br>(14–26) | 41.5<br>(30–59) |
| LVESV (mL) | 3.5<br>(2–7) | 3<br>(2–6) | 5**<br>(5–8) | 9<br>(7–15) | 5.5*<br>(4–8) | 12<br>(4–21) |
| EF (%) | 71.5<br>(58–77) | 75.5<br>(60–85) | 67<br>(57–74) | 70.5<br>(65–74) | 69<br>(55–85) | 70.5<br>(53–88) |
| **M-mode** |  |  |  |  |  |  |
| Heart Rate (bpm) | 127<br>(100–172) | 125.5<br>(94–133) | 126<br>(85–167) | 86.5<br>(66–106) | 106**<br>(82–139) | 76<br>(54–98) |
| LVIDd (cm) | 2.3<br>(2.1–3.0) | 2.4<br>(2.1–2.7) | 2.7**<br>(2.4–3.0) | 3.4<br>(3.2–4.0) | 2.9**<br>(2.8–3.3) | 3.8<br>(3.4–4.3) |
| LVIDs (cm) | 1.35<br>(1.1–1.8) | 1.3<br>(1.1–1.9) | 1.55**<br>(1.2–1.9) | 2.1<br>(1.8–2.5) | 1.6**<br>(1.3–1.9) | 2.25<br>(1.6–2.8) |
| FS (%) | 42.5<br>(38–48) | 37.5<br>(30–56) | 41.5<br>(30–54) | 40<br>(33–47) | 43.5<br>(35–56) | 42.5<br>(31–52) |
| **Strain** |  |  |  |  |  |  |
| Circ Strain (%) | -17.275<br>(-23.93 to<br>-16.15) | -17.75<br>(-23.33 to<br>-13.67) | -21.17<br>(-25.90 to<br>-13.27) | -20.17<br>(-22.83 to<br>-15.67) | -23.47<br>(-26.20 to<br>-17.27) | -20.18<br>(-24.20 to<br>-17.37) |
| Circ Strain Rate<br>(1/s) | -2.255<br>(-2.62 to<br>-1.86) | -2.28<br>(-3.04 to<br>-1.79) | -2.56*<br>(-3.33 to<br>-1.79) | -1.975<br>(-2.15 to<br>-1.56) | -2.64*<br>(-3.08 to<br>-2.01) | -1.845<br>(-2.84 to<br>-1.45) |

\* The result differed ( $P < 0.05$ ) between the affected and normal dogs.
\*\* The result differed ( $P < 0.01$ ) between the affected and normal dogs.
The complete table including carriers is shown in Table S2.

**Table 4.** CMR results from GRMD and normal dogs.

|  | 3 Months |  | 6 Months |  | 12 Months |  |
| --- | --- | --- | --- | --- | --- | --- |
|  | Affected<br>Median<br>(Min–Max)<br>N = 10 | Normal<br>Median<br>(Min–Max)<br>N = 8 | Affected<br>Median<br>(Min–Max)<br>N = 10 | Normal<br>Median<br>(Min–Max)<br>N = 8 | Affected<br>Median<br>(Min–Max)<br>N = 10 | Normal<br>Median<br>(Min–Max)<br>N = 8 |
| Heart Rate (bpm) | 127.5<br>(110–165) | 110.5<br>(98–125) | 124.5*<br>(108–148) | 105.5<br>(91–120) | 125.5**<br>(105–135) | 96<br>(84–109) |
| EF (%) | 53.05<br>(28.8–64.8) | 32.1<br>(23.1–54.1) | 47.4*<br>(28.8–54.5) | 32.35<br>(27.0–43.9) | 48.5*<br>(28.7–52.2) | 29.15<br>(21.2–45.7) |
| LVEDV (mL) | 11.75<br>(10.9–15.5) | 13.95<br>(8.9–18.6) | 23.75*<br>(16.9–28.3) | 32.65<br>(21.5–45.9) | 25.8**<br>(20.4–34.8) | 48.85<br>(40.9–67.6) |
| LVESV (mL) | 6.3<br>(3.9–8.9) | 9.45<br>(5.5–14.3) | 12.4**<br>(9.4–17.6) | 22.0<br>(12.1–33.2) | 13.95**<br>(10.1–21.5) | 35.7<br>(22.2–53.3) |
| Myo Mass (g) | 15.05 | 18.75 | 26.25** | 46.95 | 34.15** | 73.5 |
|  | Affected<br>Median<br>(Min–Max)<br>N = 10 | Normal<br>Median<br>(Min–Max)<br>N = 8 | Affected<br>Median<br>(Min–Max)<br>N = 10 | Normal<br>Median<br>(Min–Max)<br>N = 8 | Affected<br>Median<br>(Min–Max)<br>N = 10 | Normal<br>Median<br>(Min–Max)<br>N = 8 |
|  | (11.7–20.6) | (13.9–22.2) | (18.1–29.9) | (33.3–55.7) | (21.7–44.2) | (61.0–88.3) |
| SV (mL) | 5.57<br>(3.84–11.60) | 5.50<br>(3.65–6.56) | 11.695<br>(7.28–14.12) | 12.275<br>(7.65–18.09) | 12.025<br>(9.10–16.50) | 13.86<br>(12.15–20.04) |
| CO (L/min) | 0.69<br>(0.49–1.28) | 0.60<br>(0.42–0.82) | 1.39<br>(0.94–2.05) | 1.26<br>(0.83–2.17) | 1.47<br>(1.09–2.06) | 1.42<br>(1.06–1.68) |

\* The result differed ( $P < 0.05$ ) between the affected and normal dogs.
\*\* The result differed ( $P < 0.01$ ) between the affected and normal dogs.
The complete table including carriers is shown in Table S3.

#### Systolic Function and LV Outflow in Echocardiography versus CMR

Systolic function and hemodynamic results differed between the two modalities, likely because dogs were only sedated during echocardiography but anesthetized for CMR. Although the EF from CMR showed relatively lower values, the results were still comparable between genotypes since the scans were performed in a consistent manner. The EF from CMR was higher in affected dogs at 6 and 12 months (*P* < 0.05 for both) (Table 4 and S3) but did not differ at the three time points for echocardiography (Table 3 and S2).

The results of stroke volume (SV) and cardiac output (CO) differed on echocardiography but not CMR. The echocardiographic velocity time integral (VTI) at the aortic valve (AV VTI) and derived SV were lower in affected vs. normal dogs at 6 and 12 months (*P* < 0.01 for most). Moreover, affected dogs had lower cardiac output (CO) on echocardiography at 3 (*P* = 0.008) and 6 months (*P* = 0.03).

Additionally, there were incidental findings on echocardiographic analysis. Affected dogs had higher values of tricuspid valve E wave velocity (TV E Velocity) at 6 months and lower maximum velocity at the pulmonary valve (PV Vmax) at 3 months (*P* < 0.05 for both). Also, the systolic myocardial velocity at the LV lateral wall (Lateral Sm) was lower in affected dogs at 12 months (*P* = 0.03).

#### LV Diastolic Function

We did not find strong evidence of diastolic dysfunction in affected dogs. Although the ratio of the mitral valve E wave to early diastolic myocardial velocity (E/Em) at the LV lateral wall was lower in affected vs. normal dogs at 3 months (*P* = 0.04), the E/Em at both the LV lateral and septal walls did not differ between affected and normal dogs at 6 and 12 months. In assessing echocardiographic TDI results, the ratio of early to late diastolic myocardial velocity (Em/Am) at the LV septum and lateral wall was higher in affected dogs (*P* = 0.02 and 0.002) at 3 months but not at 6 and 12 months. Similarly, the ratio of mitral valve E wave to A wave (MV E/A) and the isovolumic relaxation time (IVRT) did not differ at the three time points.

#### Circumferential Strain and Strain Rate

While there were no average circumferential strain differences between the two genotypes (Table 3 and S2), the average strain rate was better (more negative) in affected dogs at 6 and 12 months (*P* = < 0.05 for both). Results of segmental circumferential strains and strain rates varied and were inconsistent between different time points (Table S2). The segmental circumferential strain at 6 months was more negative only in the anterolateral segment of the affected dogs (*P* = 0.02). For segmental circumferential strain rate, affected dogs at 6 months had better strain rate in the anterior, anterolateral (*P* < 0.01 for both), inferoseptal, and inferolateral segments (*P* < 0.05 for both), but only the inferoseptal segment differed at 12 months (*P* = 0.04).

#### CMR Myocardial Characterization

Results of segmental LGE and native T1 did not differ at any of the three time points (Figure S1 and Table S4). The LGE results were varied between normal dogs, likely because of technical challenges in blinded image assessment. The ECV values (Table S5), calculated from myocardial T1 mapping, of the segments in basal and mid-LV septum and lateral wall did not differ between two genotypes at the three time points (Figure S2). In addition, there were no differences in average ECV values (Table S5) between affected and normal dogs at the three time points.

#### Natural History of GRMD Carriers

We compared the same ECG, echocardiographic, and CMR parameters of GRMD carriers versus affected and normal dogs, finding that the carriers were generally like normal dogs, with no evidence of cardiomyopathy (Table S1-S5 and Figure S1-S2). The only differences related to gender, owing to the smaller size of females compared to males. For instance, carriers had smaller IVSd, IVSs, LA diameter, and mitral valve E wave velocity (*P* < 0.05 for all) on echocardiography and smaller LVEDV and Myo Mass on CMR (*P* < 0.05 for both) at different time points during the first year of age. Also, circumferential strain rate in the anteroseptal segment was worse (less negative) in carriers than normal dogs at 3 months (*P* = 0.04).

### Dobutamine Stress Test in the Subset Group

#### Inotropic Response of Dobutamine Stress Test

In stress echocardiography, values for EF and FS in GRMD and normal dogs did not differ at baseline prior to dobutamine dosing (Figure 4A and 4B). In response to dobutamine, EF did not differ at dosages of 5, 10, 15, and 20 µg/kg/min (Figure 4A) but was lower in affected dogs at 25 µg/kg/min (*P* = 0.03), indicating a potential decreased inotropic response. In contrast, affected dogs had lower FS values at 5, 10, 15, and 25 µg/kg/min (*P* < 0.05 for all) (Figure 4B).

Value changes (Δ) between pre-stress baseline and each dobutamine dosage were compared for GRMD and normal dogs (Figure 4C and 4D). The change in EF (ΔEF) was lower in affected vs. normal dogs at 5 (ΔEF 0-5), 10 (ΔEF 0-10), and 20 (ΔEF 0-20) µg/kg/min (*P* < 0.05 for all), no difference at 15 (ΔEF 0-15) µg/kg/min (*P* = 0.21) and showed a trend towards lowering at 25 (ΔEF 0-25) µg/kg/min (*P* = 0.053). Like ΔEF, affected dogs had lower values of FS change (ΔFS) compared to normal dogs at 5 (ΔFS 0-5), 10 (ΔFS 0-10), 15 (ΔFS 0-15), and 25 (ΔFS 0-25) µg/kg/min (*P* ≤ 0.05 for all) and a trend at 20 (ΔFS 0-20) µg/kg/min (*P* = 0.06). Taken together, ΔEF and ΔFS were all lower in affected vs. normal dogs, further documenting a decreased inotropic response.

**Figure 4.**
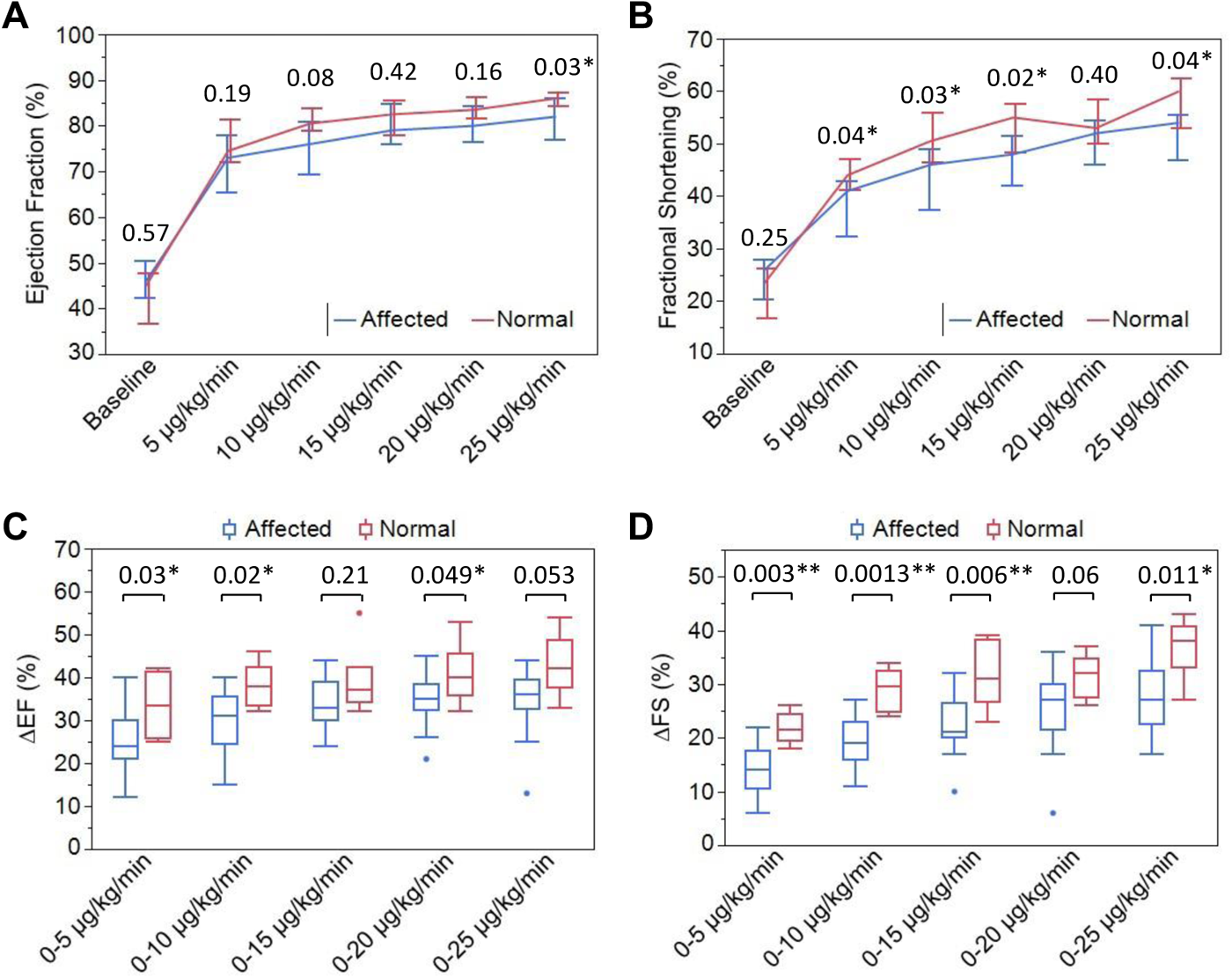
Dobutamine inotropic response in GRMD and normal dogs with echocardiography. Medians and interquartile ranges, with associated *P* values, are shown. Ejection fraction (**A**) distinguished between affected and normal dogs at 25 µg/kg/min, while fractional shortening (**B**) was more sensitive at the lower dosages. Value changes (Δ) for ejection fraction (ΔEF) (**C**) and fractional shortening (ΔFS) (**D**) also distinguished between affected and normal dogs, demonstrating a less pronounced inotropic response in affected dogs, and suggesting that ΔEF and ΔFS may be sensitive imaging markers.

In stress CMR (Figure 5A), values for EF were lower in affected vs. normal dogs at 15 (*P* = 0.03) and 20 µg/kg/min (*P* = 0.008) but not at the lower dosages. On comparison of value changes (Δ) between pre-stress baseline and each dobutamine dosage (Figure 5B), affected dogs had lower ΔEF values from baseline to 5 (ΔEF 0-5), 10 (ΔEF 0-10), 15 (ΔEF 0-15), and 20 (ΔEF 0-20) µg/kg/min (*P* < 0.05 for all), consistent with the decreased inotropic response seen in stress echocardiography.

**Figure 5.**
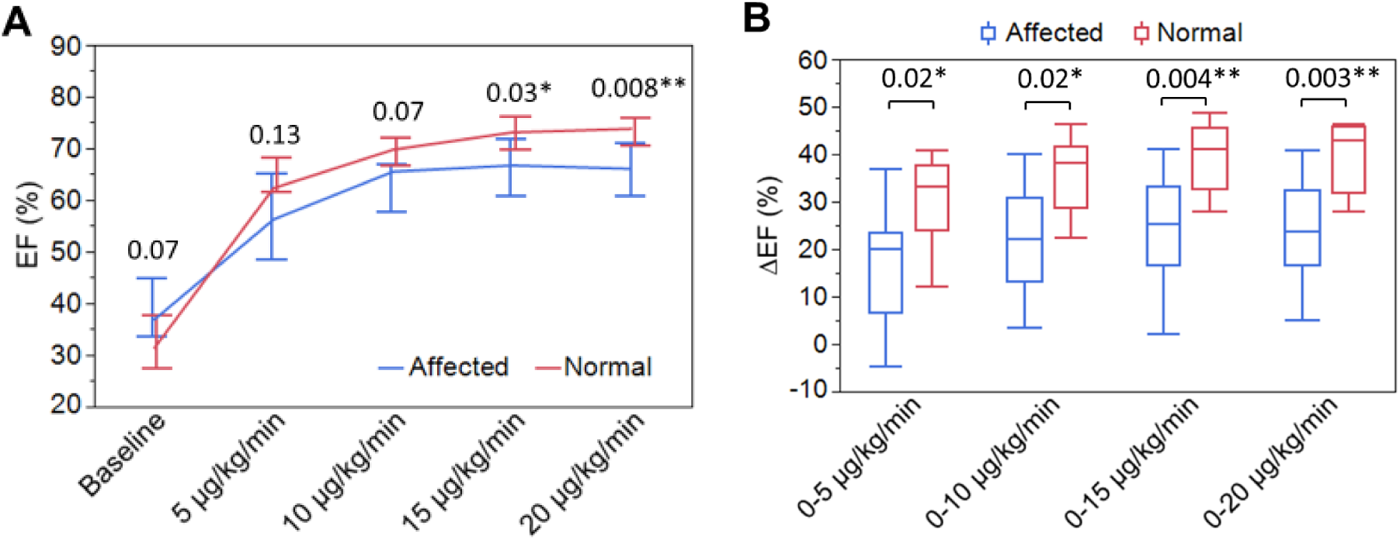
Dobutamine inotropic response in GRMD and normal dogs evaluated with CMR. Medians and interquartile ranges, with associated *P* values, are shown. Affected dogs showed lower ejection fraction values at 15 and 20 µg/kg/min (**A**). Value changes (Δ) for ejection fraction (ΔEF) were lower in affected vs. normal dogs at each dosage increment (**B**), consistent with a less pronounced inotropic response.

#### Chronotropic Response of Dobutamine Stress Test

Heart rate (HR) was compared between normal and affected dogs during stress echocardiography and CMR at 2 months of age (Figure 6A and 6B). The HR responses varied between stress echocardiography and stress CMR, even though dogs were anesthetized during both scans. Whereas HR did not differ in stress echocardiography at any dobutamine dosage, it was lower in affected dogs at 15 µg/kg/min (*P* = 0.04) during stress CMR. Value changes of heart rate (ΔHR) were also evaluated in each modality (Figure 6C and 6D). The ΔHR was lower between baseline and either 15 (ΔHR 0-15) or 20 (ΔHR 0-20) µg/kg/min (*P* < 0.05 for both) during stress CMR, but not during stress echocardiography.

**Figure 6.**
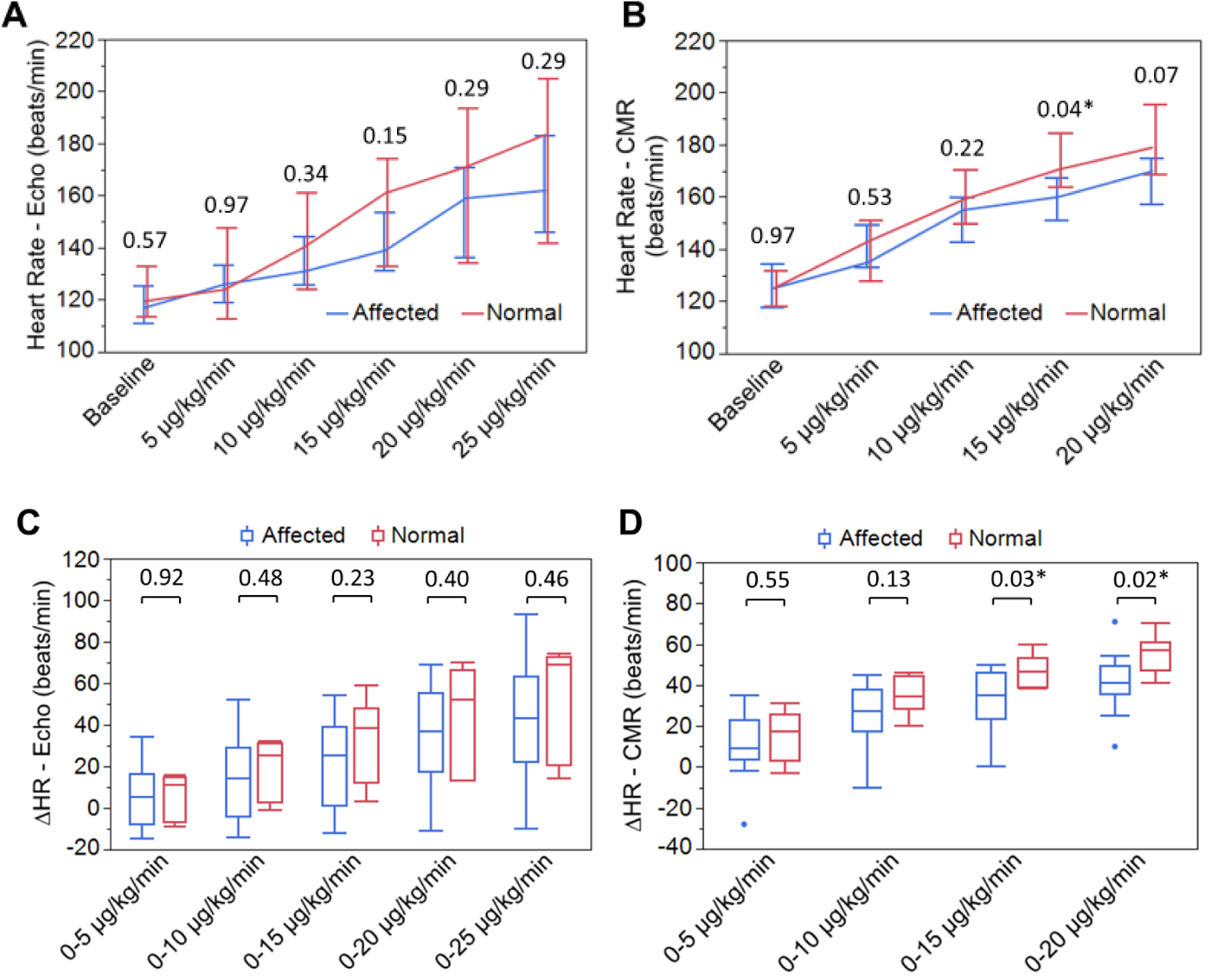
Dobutamine chronotropic response in GRMD and normal dogs with echocardiography and CMR. Medians and interquartile ranges, with associated *P* values, are shown. In stress echocardiography, heart rate did not distinguish the two genotypes (**A**). The affected dogs had lower heart rates at 15 µg/kg/min during stress CMR (**B**). The value changes of heart rate (ΔHR) in stress echocardiography (**C**) did not distinguish the two genotypes but affected dogs had lower ΔHR values at 15 and 20 µg/kg/min during stress CMR (**D**). Note that interquartile ranges are wider in affected dogs, likely reflecting phenotypic variation typical of GRMD.

#### Dobutamine Stress Test at 4.5 and 6 Months

In the follow-up stress echocardiography at 4.5 and 6 months of age, neither FS nor ΔFS distinguished the 4 affected and 6 normal dogs at dobutamine dosages of 5 and 10 µg/kg/min (Figure 7A to 7C).

**Figure 7.**
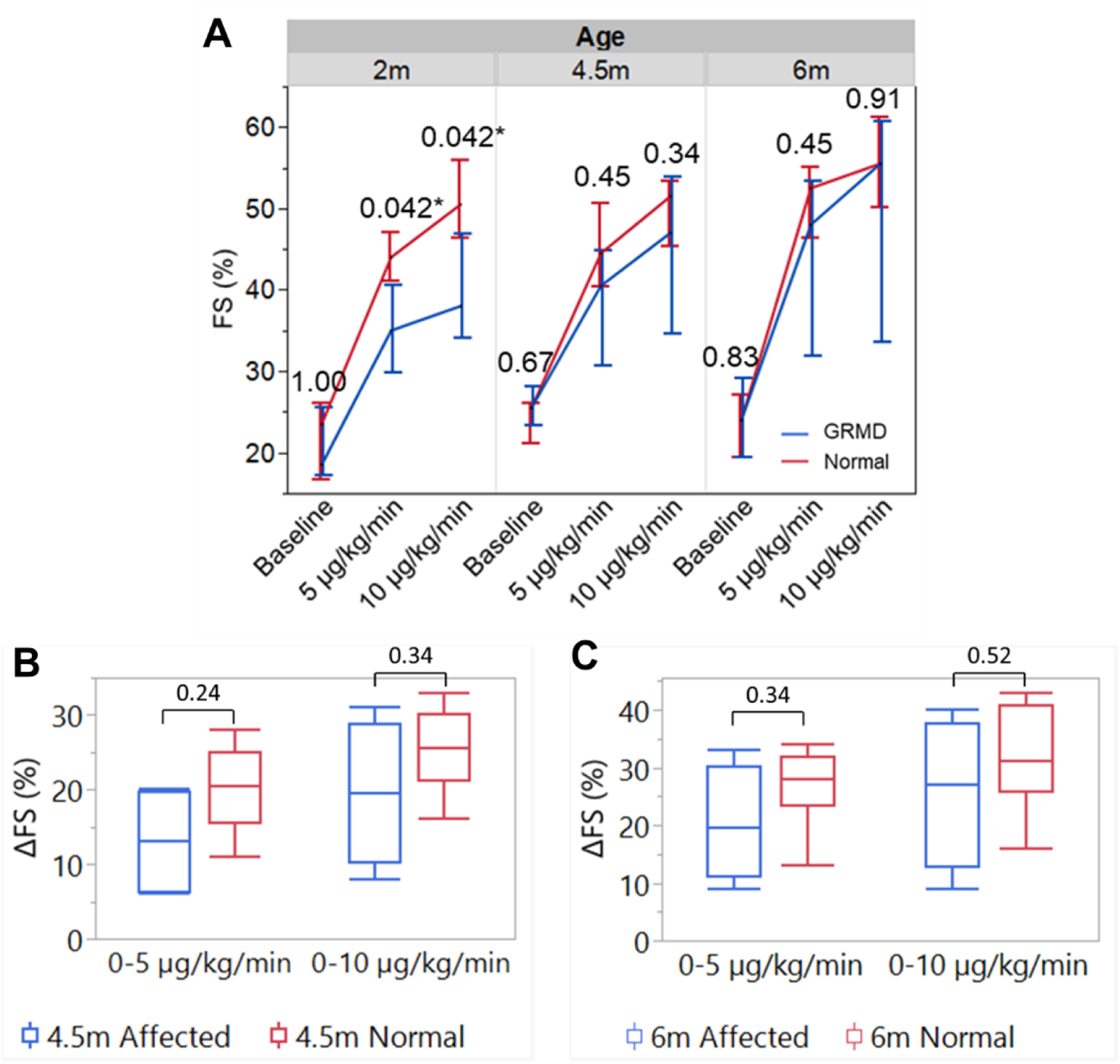
Comparisons of inotropic response between 4 affected and 6 normal dogs during dobutamine stress test. The absolute values of fractional shortening (FS) showed differences at 5and 10 µg/kg/min infusion at 2 months, but not 4.5 and 6 months (**A**). The value changes of fractional shortening (ΔFS) also did not show differences at 4.5 (**B**) and 6 months (**C**).

#### cTnI Response in Dobutamine Stress Test

Values for cTnI were compared between affected and normal dogs using three approaches. First, upon comparing values between genotypes at each pre-stress baseline and 4-hours (post-peak 4 hours) after the dobutamine peak dosage (25 µg/kg/min), there were no differences, although affected dogs showed a trend (*P* = 0.07) for higher cTnI values (Figure S3). Second, the comparison between pre- and post-peak cTnI values (Figure 8) demonstrated increased cTnI values in affected dogs at post-peak 4 hours compared to baseline (*P* = 0.02), but there was no difference in normal dogs (*P* = 0.13). Finally, change of cTnI (ΔcTnI) between pre-stress and post-peak 4 hours did not differ between the two genotypes (Figure S4).

**Figure 8.**
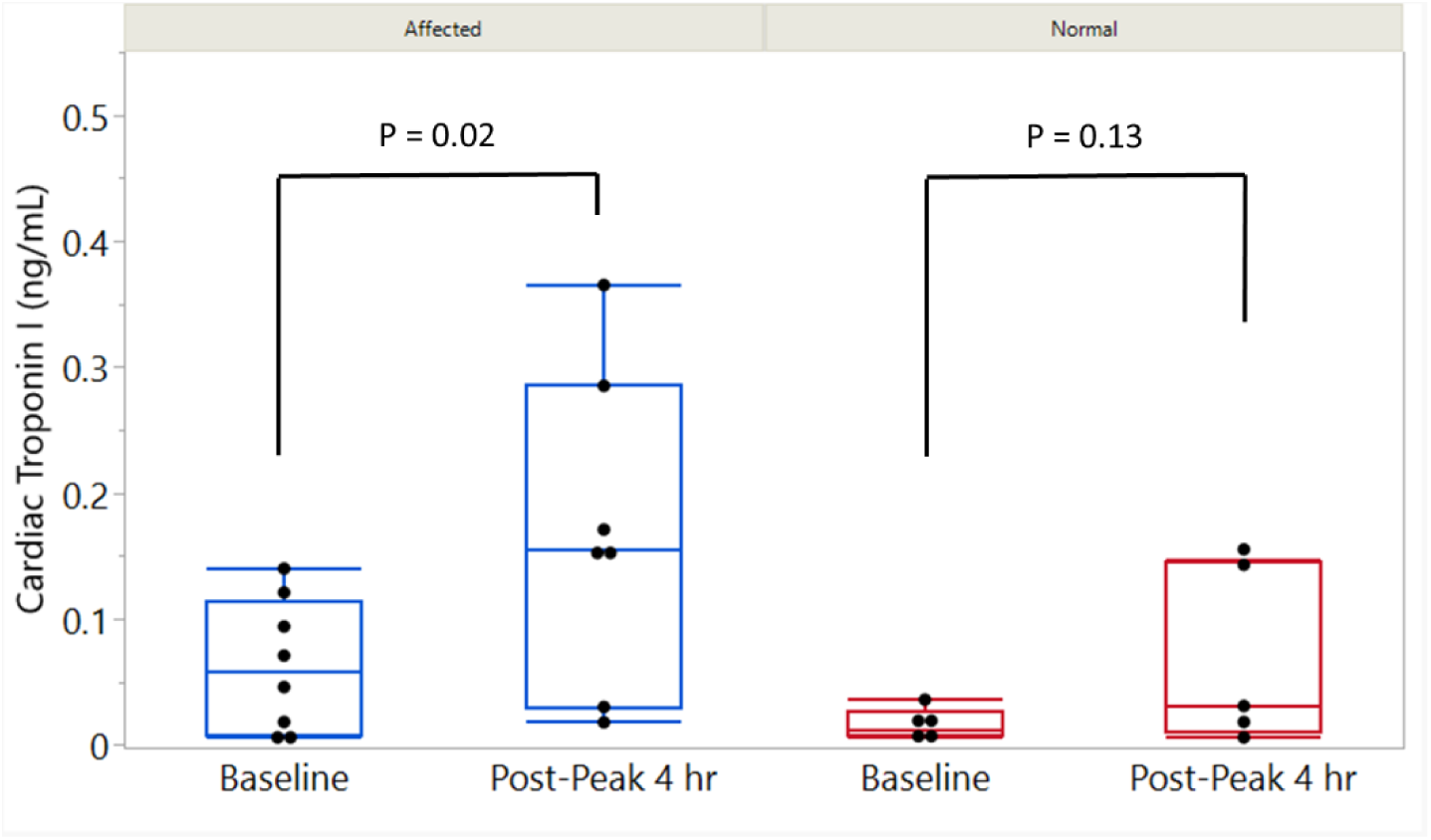
Pre- and post-stress cardiac troponin I (cTnI) values of 12 dogs (7 affected and 5 normal). One affected dog had the baseline measurement but not at post-peak 4 hours. Values for cTnI were increased between baseline and post-peak 4 hours for affected (*P* = 0.02) but not normal (*P* = 0.13) dogs.

## Discussion

Preclinical studies in GRMD are typically conducted from 3 to 6 months of age, which roughly corresponds to 5 to 10 years in DMD.^7^ While skeletal muscle disease progresses rapidly in each condition over these analogous periods, signs of cardiomyopathy occur later in the disease process for both DMD and GRMD.^9^ Accordingly, early subclinical markers are needed to assess cardiac abnormalities in juvenile GRMD dogs. In this study, we characterized the natural history of cardiomyopathy over the first year of life, using a comprehensive imaging approach. Juvenile GRMD dogs had relatively smaller hearts, characteristic ECG changes, a paradoxical phenomenon of better systolic function under anesthesia, and no clear CMR evidence of myocardial fibrosis. On the dobutamine stress test at 2 months of age, affected dogs also had a less pronounced inotropic response and increased cardiac troponin I (cTnI) between baseline and post-peak 4 hours, providing potential additional early subclinical cardiac markers.

These early changes of subclinical disease were consistent with prior GRMD studies and largely paralleled those of DMD cardiomyopathy. Abnormal ECG findings have been well documented in both DMD and GRMD.^2,25,26^ These changes precede imaging abnormalities in DMD and include sinus tachycardia, tall R waves, and increased Q wave amplitude in the left precordial leads.^2^ In studying a mixed group of juvenile and adult GRMD dogs, Moise et al found increased heart rate, deep Q waves, and increased Q/R ratio in leads II, III, aVF, V2, and V4.^25^ Similarly, Fine et al showed increased Q/R ratios in leads II, III, and aVF as early as 3 months of age in a small group of crossbred dogs with different *DMD* gene mutations.^8^ In our study, heart rate was increased in GRMD vs. normal or carrier dogs at 6 months of age but significance was not clearly established until 12 months. Consistent with the increased heart rate, the PR interval of affected dogs was shortened as early as 6 months. Like DMD, the increased heart rate and shorter PR intervals likely occurred due to the sinus tachycardia with disease progression. Therefore, the ECG changes in GRMD provide an important marker of early cardiac disease.

Our study also showed that GRMD dogs have smaller hearts compared to carrier and normal dogs starting at 6 months of age. Similar findings were found in Fine et al’s study of crossbred dogs with different *DMD* mutations beginning at this same age.^8^ An analogous less pronounced increase in percentage of cardiac chamber size at 3 to 6 months compared to normal was seen in an Australian labradoodle dystrophinopathy model.^27^ In addition, GRMD dogs in our study had less myocardial mass than carriers and normal dogs at 6 and 12 months, consistent with a smaller heart. Strikingly, GRMD myocardial masses were more than two times smaller than normal dogs by 12 months. The bodies of these affected dogs were also smaller, shown by the lower values of body weight and body surface area, compared to carriers and normal dogs suggesting that the smaller hearts could simply reflect body size. On the other hand, the 12-month GRMD dogs were only around 40% and 28% smaller than normal dogs in respective body weight and body surface area, suggesting other factors may be responsible for the disproportional myocardial mass in GRMD. Additional factors may include the inherent defect, dystrophin deficiency which leads to reduced overall activity, pathogenesis of cardiomyopathy, changes of cardiac workload, and the potential of reduced growth factors. Normalization for body size, either using body weight or BSA, should be carefully evaluated for genotype comparisons. The GRMD affected dogs had smaller body size due to the muscular dystrophy, which could bias the calculation and comparison. In any case, our study still demonstrated the smaller GRMD cardiac size could serve as a marker to differentiate genotypes.

Interestingly, GRMD dogs showed a paradoxical phenomenon of better systolic function while anesthetized for CMR. Although a differential effect of anesthesia could not be fully excluded, we suspected this was a compensatory response, together with increased heart rate, to underlying cardiac disease. This might have occurred due to higher resting sympathetic tone in GRMD dogs particularly during anesthesia. Consistent with potential autonomic dysfunction, GRMD dogs in our studies often showed progressive sinus arrhythmia and sinus tachycardia.^28^ Autonomic dysfunction also has been demonstrated in DMD boys.^29–32^ However, other parameters of systolic function, such as EF, FS, and myocardial strain on echocardiography, did not differ between genotypes while the dogs were only sedated. A somewhat analogous increase in systolic function has been seen in DMD patients with hypertrophic cardiomyopathy that subsequently progressed to the more typical dilated form.^33,34^ Regardless of the underlying pathogenesis and mechanisms, this “hyperdynamic state” associated with the increased heart rate and contractility, would be expected over time to contribute to the features of cardiomyopathy seen in older GRMD dogs,^9^ analogous to the progression of hypertrophic to dilated cardiomyopathy in DMD.^33,35^

In our previous natural history study of adult GRMD dogs,^9^ we found that changes in average circumferential strain, particularly in the anteroseptal and inferior segments, at 22 to 34 months of age were more sensitive than reduced EF in demonstrating systolic dysfunction. Consistent with these GRMD results, Ryan et al found that DMD patients less than 8 years old had worse circumferential strain at the anteroseptal, inferior, and inferolateral segments.^36^ However, the myocardial strain did not distinguish genotypes in our current study, likely because of the early disease process and relatively healthy hearts in GRMD dogs. Also in our study, myocardial strain data varied markedly in these younger dystrophic dogs even when multiple measurements were averaged, making it difficult to identify segmental abnormalities during the early disease process. Considering these inconsistencies, the few significant differences in circumferential strain rate could be related to the increased heart rates of affected dogs. Takano et al previously evaluated myocardial strains in a group of juvenile and adult CXMD_J_ (canine X-linked muscular dystrophy in Japan) beagle dogs, a canine model outbred from our GRMD colony, and showed similar non-significant findings in circumferential strain.^37^ While several studies from the French group demonstrated the potential to use longitudinal strain, cardiac twist, or MVG as early markers for GRMD dogs,^20,38^ MVG did not distinguish affected dogs in our study.

Several studies have shown that myocardial T1 mapping, with calculation of the ECV, provides a quantitative and reliable method to assess myocardial fibrosis in DMD. DMD boys were shown to have increased myocardial native T1 and ECV values compared to normal controls.^23^ Olivieri et al found similar results and further demonstrated that native T1 can identify early changes in DMD patients without the presence of LGE.^39^ Similarly, we demonstrated that adult GRMD dogs have progressively increased LGE over their lives.^9,40^ This finding is supported by a pathological study by Valentine et al in which all GRMD dogs aged ≥ 1 year had fibrosis in the subepicardial region of the LV free wall, LV papillary muscles, and right ventricular aspect of the septum,^41^ as well as a pathological study published by our group.^40^ In the current study, we used two different CMR techniques, LGE and myocardial T1 mapping, to evaluate myocardial fibrosis but did not find analogous differences across genotypes, likely reflecting mild cardiac disease early in the GRMD disease process.

Clinical cardiomyopathy occurs in DMD carriers due to skewed inactivation of their normal X-chromosome.^42,43^ We have shown that adult GRMD carriers also have pathologic cardiac changes and mosaic dystrophin expression, with associated ventricular ectopy but no reduction in systolic function.^40,44^ Largely consistent with these findings, the younger GRMD carriers studied here had no evidence of ECG abnormalities, LV dysfunction, or myocardial fibrosis, indicating a relatively healthy heart.

The GRMD colonies worldwide have shown both skeletal muscle and cardiac phenotypic differences, even though all were originally derived from the same founder GRMD dog (“Rusty”) who was diagnosed with dilated cardiomyopathy and congestive heart failure at the age of 6 years.^7,25,41^ These phenotypic differences likely reflect variable genetic backgrounds and effects of modifier genes. Over the years, dogs in our original GRMD colony have shown a milder skeletal muscle phenotype compared to those in the French and Brazilian colonies. We previously demonstrated a positive correlation between skeletal muscle and cardiac function in adult GRMD dogs,^9^ similar to DMD.^6^ Therefore, the dogs studied here may have had a relatively mild cardiac phenotype compared to what would be expected from other colonies. GRMD dogs from the French group showed LV dysfunction during the first year of life, using MVG or several advanced echocardiographic techniques.^38,45^ In any case, we did not find analogous LV dysfunction using the same techniques on our dogs.

Given our echocardiographic or CMR studies revealed few changes in young GRMD dogs, a new marker or technique other than conventional imaging methods is needed to characterize cardiac effects of early treatments. Therefore, we characterized the dobutamine stress response in GRMD by determining the ideal dosage of dobutamine and imaging markers that can distinguish affected from normal dogs by modifying published protocols used in people^14,46,47^ and dogs^5,12,48,49^ with heart disease. We found that dobutamine dosages of 5 or 10 µg/kg/min were sufficient to distinguish a less pronounced inotropic response for dystrophic dogs. A dosage up to 40 µg/kg/min increased the heart rate to approximately 170 beats per minute in most GRMD dogs without a precipitous drop of systolic function, in keeping with the target heart rate for dobutamine protocols used in people (Guo et al, unpublished data). This degree of tachycardia could increase the risk of myocardial injury and cardiac arrhythmias, as reported in a dobutamine stress test of a DMD patient.^50^

Cardiac stress tests in people are usually intended to identify regional myocardial motion abnormalities due to cardiac ischemic diseases,^14,46^ so global systolic markers are not generally evaluated. As a result, these studies are not readily applicable to DMD in which more generalized cardiac changes are expected.^2^ For the sake of animal studies, absolute EF and FS values are commonly used to evaluate the stress response.^5,48^ Importantly, because baseline values vary, absolute systolic measurements may not accurately reflect the strength of cardiac stress response. Our study showed that GRMD hearts have an apparent reduced functional response to stress, as reflected by the change of systolic markers, such as ΔEF and ΔFS, measured at pre-stress baseline and during stress testing. We compared both inotropic and chronotropic responses between GRMD and normal dogs around 2 months of age. The absolute FS and ΔFS most clearly distinguished the reduced inotropic response in GRMD dogs, while EF and ΔEF could also be used in CMR. Compared to inotropic response, the chronotropic response varied and was less sensitive in our study. Heart rate inconsistency during cardiac stress testing in animals has typically precluded assessment of the chronotropic effect.^12,48,49^ Interestingly, heart rate often temporarily decreased after starting dobutamine infusion, likely in response to vasoconstriction from the stimulation of alpha receptors.^51^

Mechanisms responsible for the reduced inotropic response in dystrophin-deficient hearts are poorly understood and likely complicated. Dobutamine exerts its inotropic effects through mixed actions on both alpha and beta myocardial receptors. Li et al showed blunting of the cardiac beta-adrenergic response in mdx mice and suggested this could be an early indicator of cardiac dysfunction in DMD.^52^ An analogous downregulation of beta receptors, with lost density, has been reported during the compensatory phase of dilated cardiomyopathy in people.^14,53,54^ As discussed, we suspected autonomic dysfunction and higher sympathetic tone in GRMD dogs as early as 6 months of age. Notably, we did not show similar blunting of the dobutamine stress response at 4.5 and 6 months of age. Therefore, other mechanisms may have been involved at 2 versus 4.5 and 6 months and, in any case, the dobutamine response test might be of lesser value at these later ages.

Serum proteins can be measured noninvasively to provide a longitudinal biomarker in DMD clinical and preclinical studies.^55^ Cardiac troponin I (cTnI) has been shown to be a sensitive serum marker of canine cardiac diseases,^56,57^ including dilated cardiomyopathy.^58^ Voleti et al also showed cTnI was elevated in DMD patients with myocardial lesions, providing a marker to monitor DMD cardiomyopathy.^59^ We hypothesized that the dystrophic heart would more likely release cTnI into serum, particularly in response to dobutamine and found affected dogs showed a significant increase of cTnI levels between pre-stress baseline and post-peak 4-hour time points and a trend towards higher cTnI levels at the post-peak 4-hour time point compared to the normal dogs. Therefore, like DMD, cTnI may be a potential serum biomarker for GRMD cardiomyopathy.

Although our studies included the largest set of young GRMD dogs yet characterized by advanced cardiac imaging, there were still limitations. In keeping with skeletal muscle disease, the cardiac phenotype varies among GRMD dogs, making it difficult to reach statistical significance. To acquire enough dogs, these studies extended across several years, during which time our experience in echocardiography and CMR improved. Also, several new imaging techniques, such as myocardial T1 mapping, have continued to advance and could potentially have been more sensitive in detecting cardiac disease. Correlative pathologic evaluations were precluded because these dogs were transferred to other projects at the end of the study. With that said, our group has recently published an extensive natural history pathologic sturdy of GRMD cardiomyopathy.^40^ In the cardiac stress study, the small hearts of 2-month-old puppies led to technical difficulties in assessing scans, particularly in making measurements at the end-systolic stage. Image quality with both echocardiography and CMR was decreased because of the increased heart rate during dobutamine stress. Although we separated the stress echocardiography and stress CMR procedures by at least 24 hours, the CMR findings could still potentially have been influenced by the stress test in echocardiography. Regarding the cTnI experiments, interpretation was compounded by the fact that the time of serum sample collection for the post-peak 4-hour time point varied by ± 1 hour. Finally, the smaller sample size at 4.5 and 6 months in the study undoubtedly made it more difficult to reach statistical significance.

## Conclusions

This study characterized the natural history of cardiomyopathy over the first year of life in GRMD dogs compared to age-matched normal and carrier dogs using ECG, echocardiography, and CMR. We found that GRMD dogs had early ECG changes, showing higher heart rates, increased Q/R ratios, and shortening of the PR interval. They also had smaller hearts and disproportionally less myocardial mass at 6 and 12 months of age. No systolic dysfunction or consistent differences in circumferential strain across genotypes were found, but GRMD dogs showed a paradoxical phenomenon of better systolic function under anesthesia during CMR. Although myocardial fibrosis was not found using LGE and myocardial T1 mapping, these techniques should still be helpful in future studies. Findings from GRMD carriers were comparable to those from normal dogs, with no evidence of cardiomyopathy at this early age. In addition, we established a dobutamine protocol whereby dosages as low as 5 or 10 µg/kg/min could be used to differentiate GRMD vs. normal dogs at 2 months of age. Combined with our previous study of adult GRMD dogs, our findings provide the most comprehensive natural history cardiac imaging data for young and adult GRMD dogs. The remarkable parallels between the natural histories of cardiomyopathy in dystrophic dogs and boys further validate GRMD as an important model of DMD cardiomyopathy.

## Acknowledgements

We acknowledge Dr. Alan C. Glowczwski for the imaging support of the CMR scans and Comparative Medicine Program at Texas A&M University for the care of the study animals. In addition, we would like to acknowledge Dr. Paul T. Martin and Solid Biosciences for letting us collect the cardiac stress data prior to their studies.

## Author Contributions

Guo performed the main study and image processing, analyzed data, and wrote the manuscript. Guo performed all the ECG and echocardiographic scans in this study. Bettis assisted with animal procedures. Nghiem, Soslow, Spurney, and Kornegay assisted with the analytic approach and manuscript preparation. Kornegay provided funding for the study and supervised the overall study. All authors participated in revision of the manuscript.

## References

1. Hoffman EP, Brown RH, Jr., Kunkel LM. Dystrophin: the protein product of the Duchenne muscular dystrophy locus. Cell. 1987;51:919–928.

2. Kamdar F, Garry DJ. Dystrophin-deficient cardiomyopathy. J Am Coll Cardiol. 2016;67:2533–2546. doi: 10.1016/j.jacc.2016.02.081

3. Marty B, Gilles R, Toussaint M, Behin A, Stojkovic T, Eymard B, Carlier PG, Wahbi K. Comprehensive evaluation of structural and functional myocardial impairments in Becker muscular dystrophy using quantitative cardiac magnetic resonance imaging. Eur Heart J Cardiovasc Imaging. 2019;20:906–915. doi: 10.1093/ehjci/jey209

4. Birch SM, Lawlor MW, Conlon TJ, Guo LJ, Crudele JM, Hawkins EC, Nghiem PP, Ahn M, Meng H, Beatka MJ, et al. Assessment of systemic AAV-microdystrophin gene therapy in the GRMD model of Duchenne muscular dystrophy. Sci Transl Med. 2023;15:eabo1815. doi: 10.1126/scitranslmed.abo1815

5. Townsend D, Turner I, Yasuda S, Martindale J, Davis J, Shillingford M, Kornegay JN, Metzger JM. Chronic administration of membrane sealant prevents severe cardiac injury and ventricular dilatation in dystrophic dogs. J Clin Invest. 2010;120:1140–1150. doi: 10.1172/jci41329

6. Posner AD, Soslow JH, Burnette WB, Bian A, Shintani A, Sawyer DB, Markham LW. The Correlation of Skeletal and Cardiac Muscle Dysfunction in Duchenne Muscular Dystrophy. J Neuromuscul Dis. 2016;3:91–99. doi: 10.3233/jnd-150132

7. Kornegay JN. The golden retriever model of Duchenne muscular dystrophy. Skelet Muscle. 2017;7:9. doi: 10.1186/s13395-017-0124-z

8. Fine DM, Shin JH, Yue Y, Volkmann D, Leach SB, Smith BF, McIntosh M, Duan D. Age-matched comparison reveals early electrocardiography and echocardiography changes in dystrophin-deficient dogs. Neuromuscul Disord. 2011;21:453–461. doi: 10.1016/j.nmd.2011.03.010

9. Guo LJ, Soslow JH, Bettis AK, Nghiem PP, Cummings KJ, Lenox MW, Miller MW, Kornegay JN, Spurney CF. Natural History of Cardiomyopathy in Adult Dogs With Golden Retriever Muscular Dystrophy. Journal of the American Heart Association. 2019;8:e012443. doi: 10.1161/jaha.119.012443

10. Hammers DW, Sleeper MM, Forbes SC, Shima A, Walter GA, Sweeney HL. Tadalafil Treatment Delays the Onset of Cardiomyopathy in Dystrophin-Deficient Hearts. Journal of the American Heart Association. 2016;5. doi: 10.1161/jaha.116.003911

11. Li Z, Li Y, Zhang L, Zhang X, Sullivan R, Ai X, Szeto C, Cai A, Liu L, Xiao W, et al. Reduced Myocardial Reserve in Young X-Linked Muscular Dystrophy Mice Diagnosed by Two-Dimensional Strain Analysis Combined with Stress Echocardiography. J Am Soc Echocardiogr. 2017;30:815–827.e819. doi: 10.1016/j.echo.2017.03.009

12. McEntee K, Amory H, Pypendop B, Balligand M, Clercx C, Michaux C, Jacqmot O, Robert F, Gerard P, Pochet T, et al. Effects of dobutamine on isovolumic and ejection phase indices of cardiac contractility in conscious healthy dogs. Res Vet Sci. 1998;64:45–50.

13. Maruo T, Nakatani S, Jin Y, Uemura K, Sugimachi M, Ueda-Ishibashi H, Kitakaze M, Ohe T, Sunagawa K, Miyatake K. Evaluation of transmural distribution of viable muscle by myocardial strain profile and dobutamine stress echocardiography. Am J Physiol Heart Circ Physiol. 2007;292:H921–927. doi: 10.1152/ajpheart.00019.2006

14. Lancellotti P, Pellikka PA, Budts W, Chaudhry FA, Donal E, Dulgheru R, Edvardsen T, Garbi M, Ha JW, Kane GC, et al. The Clinical Use of Stress Echocardiography in Non-Ischaemic Heart Disease: Recommendations from the European Association of Cardiovascular Imaging and the American Society of Echocardiography. J Am Soc Echocardiogr. 2017;30:101–138. doi: 10.1016/j.echo.2016.10.016

15. Bartlett RJ, Winand NJ, Secore SL, Singer JT, Fletcher S, Wilton S, Bogan DJ, Metcalf-Bogan JR, Bartlett WT, Howell JM, et al. Mutation segregation and rapid carrier detection of X-linked muscular dystrophy in dogs. Am J Vet Res. 1996;57:650–654.

16. Kornegay JN, Bogan JR, Bogan DJ, Childers MK, Li J, Nghiem P, Detwiler DA, Larsen CA, Grange RW, Bhavaraju-Sanka RK, et al. Canine models of Duchenne muscular dystrophy and their use in therapeutic strategies. Mamm Genome. 2012;23:85–108. doi: 10.1007/s00335-011-9382-y

17. 17. Guo LJ, Spurney CF. Large mammal (canine) ECG. Parent Project Muscular Dystrophy. https://www.parentprojectmd.org/wp-content/uploads/2018/04/3.1.Dog_ECG_42815.DF_.DDa_.pdf. 2015

18. Wess G, Maurer J, Simak J, Hartmann K. Use of Simpson’s method of disc to detect early echocardiographic changes in Doberman Pinschers with dilated cardiomyopathy. J Vet Intern Med. 2010;24:1069–1076. doi: 10.1111/j.1939-1676.2010.0575.x

19. Spurney CF, McCaffrey FM, Cnaan A, Morgenroth LP, Ghelani SJ, Gordish-Dressman H, Arrieta A, Connolly AM, Lotze TE, McDonald CM, et al. Feasibility and Reproducibility of Echocardiographic Measures in Children with Muscular Dystrophies. J Am Soc Echocardiogr. 2015;28:999–1008. doi: 10.1016/j.echo.2015.03.003

20. Chetboul V, Escriou C, Tessier D, Richard V, Pouchelon JL, Thibault H, Lallemand F, Thuillez C, Blot S, Derumeaux G. Tissue Doppler imaging detects early asymptomatic myocardial abnormalities in a dog model of Duchenne’s cardiomyopathy. Eur Heart J. 2004;25:1934–1939. doi: 10.1016/j.ehj.2004.09.007

21. Su JB, Cazorla O, Blot S, Blanchard-Gutton N, Ait Mou Y, Barthelemy I, Sambin L, Sampedrano CC, Gouni V, Unterfinger Y, et al. Bradykinin restores left ventricular function, sarcomeric protein phosphorylation, and e/nNOS levels in dogs with Duchenne muscular dystrophy cardiomyopathy. Cardiovasc Res. 2012;95:86–96. doi: 10.1093/cvr/cvs161

22. Haaf P, Garg P, Messroghli DR, Broadbent DA, Greenwood JP, Plein S. Cardiac T1 Mapping and Extracellular Volume (ECV) in clinical practice: a comprehensive review. J Cardiovasc Magn Reson. 2016;18:89. doi: 10.1186/s12968-016-0308-4

23. Soslow JH, Damon SM, Crum K, Lawson MA, Slaughter JC, Xu M, Arai AE, Sawyer DB, Parra DA, Damon BM, et al. Increased myocardial native T1 and extracellular volume in patients with Duchenne muscular dystrophy. J Cardiovasc Magn Reson. 2016;18:5. doi: 10.1186/s12968-016-0224-7

24. 24. Plumb DC. Plumb’s veterinary drug handbook. Pocket 8th edition ed. Chichester, West Sussex: Wiley Blackwell; 2015.

25. Moise NS, Valentine BA, Brown CA, Erb HN, Beck KA, Cooper BJ, Gilmour RF. Duchenne’s cardiomyopathy in a canine model: electrocardiographic and echocardiographic studies. J Am Coll Cardiol. 1991;17:812–820.

26. Barthelemy I, Su JB, Cauchois X, Relaix F, Ghaleh B, Blot S. Ambulatory electrocardiographic longitudinal monitoring in a canine model for Duchenne muscular dystrophy identifies decreased very low frequency power as a hallmark of impaired heart rate variability. Sci Rep. 2024;14:8969. doi: 10.1038/s41598-024-59196-z

27. Shrader SM, Jung S, Denney TS, Smith BF. Characterization of Australian Labradoodle dystrophinopathy. Neuromuscul Disord. 2018;28:927–937. doi: 10.1016/j.nmd.2018.08.008

28. Rutledge AM, Guo LJ, Lord LE, Leal AR, Deramus J, Lopez SM, Russell A, Nghiem PP. Comprehensive assessment of physical activity correlated with muscle function in canine Duchenne muscular dystrophy. Ann Phys Rehabil Med. 2022;65:101611. doi: 10.1016/j.rehab.2021.101611

29. Inoue M, Mori K, Hayabuchi Y, Tatara K, Kagami S. Autonomic function in patients with Duchenne muscular dystrophy. Pediatr Int. 2009;51:33–40. doi: 10.1111/j.1442-200X.2008.02656.x

30. Thomas TO, Jefferies JL, Lorts A, Anderson JB, Gao Z, Benson DW, Hor KN, Cripe LH, Urbina EM. Autonomic dysfunction: a driving force for myocardial fibrosis in young Duchenne muscular dystrophy patients? Pediatr Cardiol. 2015;36:561–568. doi: 10.1007/s00246-014-1050-z

31. da Silva TD, Massetti T, Crocetta TB, de Mello Monteiro CB, Carll A, Vanderlei LCM, Arbaugh C, Oliveira FR, de Abreu LC, Ferreira Filho C, et al. Heart Rate Variability and Cardiopulmonary Dysfunction in Patients with Duchenne Muscular Dystrophy: A Systematic Review. Pediatr Cardiol. 2018;39:869–883. doi: 10.1007/s00246-018-1881-0

32. Lanza GA, Dello Russo A, Giglio V, De Luca L, Messano L, Santini C, Ricci E, Damiani A, Fumagalli G, De Martino G, et al. Impairment of cardiac autonomic function in patients with Duchenne muscular dystrophy: relationship to myocardial and respiratory function. Am Heart J. 2001;141:808–812. doi: 10.1067/mhj.2001.114804

33. Nigro G, Comi LI, Politano L, Bain RJ. The incidence and evolution of cardiomyopathy in Duchenne muscular dystrophy. Int J Cardiol. 1990;26:271–277.

34. Canonico F, Chirivi M, Maiullari F, Milan M, Rizzi R, Arcudi A, Galli M, Pane M, Gowran A, Pompilio G, et al. Focus on the road to modelling cardiomyopathy in muscular dystrophy. Cardiovasc Res. 2022;118:1872–1884. doi: 10.1093/cvr/cvab232

35. D’Amario D, Amodeo A, Adorisio R, Tiziano FD, Leone AM, Perri G, Bruno P, Massetti M, Ferlini A, Pane M, et al. A current approach to heart failure in Duchenne muscular dystrophy. Heart. 2017;103:1770–1779. doi: 10.1136/heartjnl-2017-311269

36. Ryan TD, Taylor MD, Mazur W, Cripe LH, Pratt J, King EC, Lao K, Grenier MA, Jefferies JL, Benson DW, et al. Abnormal circumferential strain is present in young Duchenne muscular dystrophy patients. Pediatr Cardiol. 2013;34:1159–1165. doi: 10.1007/s00246-012-0622-z

37. Takano H, Fujii Y, Yugeta N, Takeda S, Wakao Y. Assessment of left ventricular regional function in affected and carrier dogs with Duchenne muscular dystrophy using speckle tracking echocardiography. BMC Cardiovasc Disord. 2011;11:23. doi: 10.1186/1471-2261-11-23

38. Ghaleh B, Barthélemy I, Sambin L, Bizé A, Hittinger L, Blot S, Su JB. Alteration in Left Ventricular Contractile Function Develops in Puppies With Duchenne Muscular Dystrophy. J Am Soc Echocardiogr. 2020;33:120–129.e121. doi: 10.1016/j.echo.2019.08.003

39. Olivieri LJ, Kellman P, McCarter RJ, Cross RR, Hansen MS, Spurney CF. Native T1 values identify myocardial changes and stratify disease severity in patients with Duchenne muscular dystrophy. J Cardiovasc Magn Reson. 2016;18:72. doi: 10.1186/s12968-016-0292-8

40. Schneider SM, Sansom GT, Guo LJ, Furuya S, Weeks BR, Kornegay JN. Natural History of Histopathologic Changes in Cardiomyopathy of Golden Retriever Muscular Dystrophy. Front Vet Sci. 2021;8:759585. doi: 10.3389/fvets.2021.759585

41. Valentine BA, Cummings JF, Cooper BJ. Development of Duchenne-type cardiomyopathy. Morphologic studies in a canine model. Am J Pathol. 1989;135:671–678.

42. Soltanzadeh P, Friez MJ, Dunn D, von Niederhausern A, Gurvich OL, Swoboda KJ, Sampson JB, Pestronk A, Connolly AM, Florence JM, et al. Clinical and genetic characterization of manifesting carriers of DMD mutations. Neuromuscul Disord. 2010;20:499–504. doi: 10.1016/j.nmd.2010.05.010

43. Lim KRQ, Sheri N, Nguyen Q, Yokota T. Cardiac Involvement in Dystrophin-Deficient Females: Current Understanding and Implications for the Treatment of Dystrophinopathies. Genes (Basel). 2020;11. doi: 10.3390/genes11070765

44. Kane AM, DeFrancesco TC, Boyle MC, Malarkey DE, Ritchey JW, Atkins CE, Cullen JM, Kornegay JN, Keene BW. Cardiac structure and function in female carriers of a canine model of Duchenne muscular dystrophy. Res Vet Sci. 2013;94:610–617. doi: 10.1016/j.rvsc.2012.09.027

45. Ghaleh B, Barthélemy I, Wojcik J, Sambin L, Bizé A, Hittinger L, Tran TD, Thomé FP, Blot S, Su JB. Protective effects of rimeporide on left ventricular function in golden retriever muscular dystrophy dogs. Int J Cardiol. 2020;312:89–95. doi: 10.1016/j.ijcard.2020.03.031

46. Armstrong WF, Zoghbi WA. Stress echocardiography: current methodology and clinical applications. J Am Coll Cardiol. 2005;45:1739–1747. doi: 10.1016/j.jacc.2004.12.078

47. Nabi F, Malaty A, Shah DJ. Stress cardiac magnetic resonance. Curr Opin Cardiol. 2011;26:385–391. doi: 10.1097/HCO.0b013e3283490373

48. Suzuki R, Matsumoto H, Teshima T, Mochizuki Y, Koyama H. Dobutamine stress echocardiography for assessment of systolic function in dogs with experimentally induced mitral regurgitation. J Vet Intern Med. 2014;28:386–392. doi: 10.1111/jvim.12293

49. Spasojevic Kosic L, Trailovic DR, Matunovic R. Resting and dobutamine stress test induced serum concentrations of brain natriuretic peptide in German Shepherd dogs. Res Vet Sci. 2012;93:1446–1453. doi: 10.1016/j.rvsc.2012.04.002

50. Wong BL, Mukkada VA, Markham LW, Cripe LH. Depressed left ventricular contractile reserve diagnosed by dobutamine stress echocardiography in a patient with Duchenne muscular dystrophy. J Child Neurol. 2005;20:246–248. doi: 10.1177/08830738050200031401

51. Bistola V, Chioncel O. Inotropes in acute heart failure. Continuing Cardiology Education. 2017;3:107–116. doi: 10.1002/cce2.59

52. Li Y, Zhang S, Zhang X, Li J, Ai X, Zhang L, Yu D, Ge S, Peng Y, Chen X. Blunted cardiac beta-adrenergic response as an early indication of cardiac dysfunction in Duchenne muscular dystrophy. Cardiovasc Res. 2014;103:60–71. doi: 10.1093/cvr/cvu119

53. Wilkinson R, Song W, Smoktunowicz N, Marston S. A dilated cardiomyopathy mutation blunts adrenergic response and induces contractile dysfunction under chronic angiotensin II stress. Am J Physiol Heart Circ Physiol. 2015;309:H1936–1946. doi: 10.1152/ajpheart.00327.2015

54. Ruffolo RR, Jr. The pharmacology of dobutamine. Am J Med Sci. 1987;294:244–248. doi: 10.1097/00000441-198710000-00005

55. Hathout Y, Brody E, Clemens PR, Cripe L, DeLisle RK, Furlong P, Gordish-Dressman H, Hache L, Henricson E, Hoffman EP, et al. Large-scale serum protein biomarker discovery in Duchenne muscular dystrophy. Proc Natl Acad Sci U S A. 2015;112:7153–7158. doi: 10.1073/pnas.1507719112

56. Spratt DP, Mellanby RJ, Drury N, Archer J. Cardiac troponin I: evaluation I of a biomarker for the diagnosis of heart disease in the dog. J Small Anim Pract. 2005;46:139–145. doi: 10.1111/j.1748-5827.2005.tb00304.x

57. Langhorn R, Willesen JL. Cardiac Troponins in Dogs and Cats. J Vet Intern Med. 2016;30:36–50. doi: 10.1111/jvim.13801

58. Wess G, Simak J, Mahling M, Hartmann K. Cardiac troponin I in Doberman Pinschers with cardiomyopathy. J Vet Intern Med. 2010;24:843–849. doi: 10.1111/j.1939-1676.2010.0516.x

59. Voleti S, Olivieri L, Hamann K, Gordish-Dressman H, Spurney C. Troponin I Levels Correlate with Cardiac MR LGE and Native T1 Values in Duchenne Muscular Dystrophy Cardiomyopathy and Identify Early Disease Progression. Pediatr Cardiol. 2020;41:1173–1179. doi: 10.1007/s00246-020-02372-5

